# A Simple, Cost-Effective, High-Throughput Method for Measuring Chromatin Accessibility and Gene Expression in Single Nuclei

**DOI:** 10.64898/2026.06.29.735326

**Authors:** Zhifei Luo, William J Greenleaf

## Abstract

We describe microfluidic-free, droplet-based methods for single-nucleus epigenomic measurements: Particle-templated Instant Partition single-nucleus assay for transposase-accessible chromatin using sequencing (PIP-ATAC-seq) and its multiomic version (PIP-Multiome-seq). We benchmarked these assays by generating data sets containing thousands of nuclei using cell lines and mouse brains and compared to other established methods. PIP-Multiome and PIP-ATAC are straightforward to implement, affordable, and produce high-quality data, providing useful additions to the single-cell molecular measurement armamentarium.

## Main

Chromatin accessibility is associated with “active” regulatory elements that drive gene expression programs that define cellular identity. Single-nucleus chromatin accessibility mapping in complex tissues has allowed a mapping of gene regulatory mechanisms across neuroscience, development, disease, and aging^1–3^. Yet widespread adoption of single cell chromatin accessibility methods has been hampered by the limitations of existing single-cell methods, which are often prohibitively expensive or technically demanding with inconsistent results.

Current approaches for mapping chromatin accessibility — with or without co-profiling of gene expression — fall into three categories, each with notable drawbacks. Plate-based methods require sorting individual cells into wells from which sequencing libraries are generated, making them low-throughput and costly. Microfluidics-based methods depend on specialized, expensive equipment and reagents to encapsulate single cells in droplets. Finally, split-pool-based methods are technically cumbersome, subjecting nuclei to multiple rounds of time-intensive reactions that increase ambient RNA contamination, promote nuclei clumping, and ultimately compromise data quality. A simpler, more accessible, and cost-effective alternative to these approaches would likely enable a substantially broader deployment of single-cell chromatin characterization at the single cell level^4,5^.

To address these limitations, we developed a novel single-nucleus chromatin accessibility method built on Particle-templated Instant Partitioning Sequencing (PIP-seq), an emerging alternative platform for single-cell assays^4^. In PIP-seq, uniformly sized beads serve as physical templates to generate monodispersed, water-in-oil droplets within minutes through simple vortexing — eliminating the need for specialized microfluidic hardware. Nuclei and barcoded beads are co-encapsulated within the same droplets, enabling straightforward single-nucleus library preparation directly from the partitioned emulsion.

Relative to existing approaches, our method offers several key advantages: it is substantially more cost-effective than microfluidics-based platforms, considerably higher in throughput than plate-based methods, and simpler to execute than split-pool strategies, with reduced nuclei clumping and minimal nucleic acid leakage. The method supports profiling of chromatin accessibility alone or in combination with the transcriptome. Using an adapted commercially available kit, this approach can generate data within two days without costly specialized equipment, while reliably producing high-complexity sequencing libraries — making it broadly accessible to laboratories without extensive single-cell genomics infrastructure.

We first developed PIP-ATAC-seq, which is carried out in the following steps: (1) Insertion of sequencing adapters into open chromatin regions within permeabilized nuclei using hyperactive Tn5 transposase; (2) Compartmentalization of individual nuclei together with a single barcoded bead via particle-templated instant partitioning; (3) Cell lysis through fusion of microemulsions containing lysis reagent (4) Release of chromatin fragments from the nuclei and hybridization to capture oligos on barcoded beads through splint oligos; (4) Ligation of chromatin fragments onto the beads; (5) Amplification of sequencing libraries from the beads (Fig. 1A, Fig. S1).

**Fig. 1.**
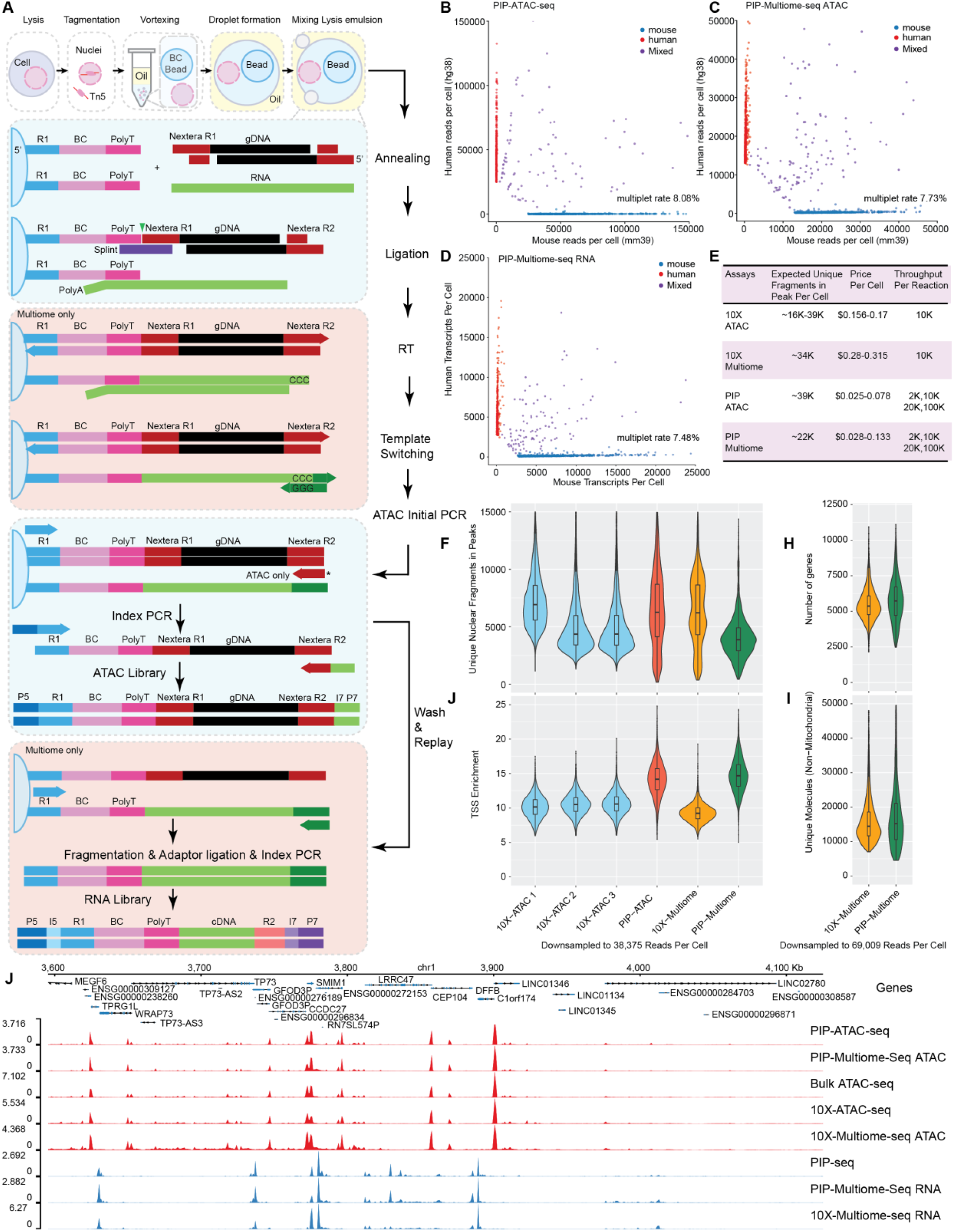
Measuring single-cell chromatin accessibility and gene expression using PIP-ATAC-seq and PIP-Multiome-seq. **a**, Overview of the PIP-ATAC-seq and PIP-Multiome-seq workflows. The two methods share most steps; PIP-ATAC-seq can be extended to multiome by incorporating additional multiome-specific steps (red box). The ATAC primer marked with * is used exclusively in PIP-ATAC-seq. **b–d**, Species mixing test for PIP-ATAC-seq (**b**), PIP-Multiome-seq ATAC (**c**), and PIP-Multiome-seq RNA (**d**). **e**, Benchmarking of PIP-ATAC-seq and PIP-Multiome-seq. Expected unique fragments per cell are based on simulated results at infinite sequencing depth. The fraction of reads in peaks was calculated on a downsampled dataset using 1,615 cells. Pricing for PIP-seq is based on quotes for the T2 to T100 kit range; pricing for 10x is based on quotes for 4- and 16-reaction kits. **f–g**, ATAC-seq benchmarking: median unique nuclear fragments per cell (**f**) and TSS enrichment (**g**). **h–i**, RNA-seq benchmarking: number of transcripts (**h**) and genes detected (**i**) per cell. **j**, Genome browser tracks for cross-method comparison.

We next extended the method to enable simultaneous profiling of chromatin accessibility and gene expression from the same nucleus. The PIP-multiome-seq workflow shares the same step (1) to (3) as PIP-ATAC-seq with the following differences: Upon lysis in step (4), released chromatin fragments and RNA are captured by oligonucleotides on the barcoded bead — chromatin fragments via ligation and RNA via reverse transcription into cDNA — while ATAC-seq fragments are simultaneously extended. ATAC-seq fragments then undergo linear amplification, are eluted from the beads, and are indexed by PCR. The beads are then washed and used for whole-transcriptome amplification, after which cDNA is purified, fragmented, ligated with sequencing adapters, and indexed by PCR.

This sequential processing of chromatin and RNA from the same barcoded bead enables paired single-nucleus accessibility and transcriptomic profiling (Fig. 1A, Fig. S1).

As proof of concept, we tested PIP-ATAC-seq using the human K562 cell line. We targeted 2,000 cells and sequenced our ATAC-seq fragment library such that the number of reads was approximately double the number of aligned fragments. Based on read complexity per cell, we identified 1,800 cells and then applied quality control filtering (Fig. S3A–E). Of these, 1,682 cells (93.4%) passed quality control, with a median of 41,801 fragments per cell and a median TSS enrichment score of 14.3. The insert size distribution exhibited clear nucleosome periodicity typical of ATAC-seq data ^6^ (Fig. S3 C).

To assess single-cell specificity, we performed a barnyard experiment by mixing human K562 and mouse 802T4 cells prior to library preparation. We recovered 750 mouse cells (median 45,923 fragments; TSS enrichment score = 23.5) and 271 human cells (median 44,435 fragments; TSS enrichment score = 15.4), indicating robust library complexity and data quality for both species. Cross-contamination was minimal, with only 0.27% of human fragments detected in mouse cells and 0.42% of mouse fragments in human cells, confirming high single-cell purity. The observed multiplet rate was 8.08% (97/1,200), similar to the <8% rate specified by the PIP-seq kit; this rate can be reduced by lowering nuclei input (Fig. 1 B, Fig. S3 G).

We also validated PIP-multi-seq using frozen K562 cells. RNA libraries contain 1,717 cells based on elbow plot (Fig. S3 B). ATAC libraries were sequenced to 52.8% saturation, and 94.1% of cells passing the RNA filter also passed the ATAC filter. From the final 1,615 high-quality cells, we detected a median of 15,428 transcripts (5,740 genes) and 25,958 ATAC fragments per cell, with a median TSS enrichment score of 15.2 (Fig. S3 D-F).

To evaluate cross-contamination for the multiomic version, we performed a barnyard experiment mixing human K562 and mouse 802T4 cells. We recovered 556 human and 835 mouse cells with a multiplet rate of 7.5%. Cross-species RNA contamination was low (2.38% mouse transcripts in human cells; 2.02% in the reverse direction), and 99% of cells passing RNA filters also passed ATAC filters. ATAC-seq yielded 832 mouse cells (median 18,681 fragments; TSS = 24.2) and 553 human cells (median 18,257 fragments; TSS = 15.3), with cross-species fragment contamination of only 1.61% and 1.25%, respectively, confirming high single-nucleus purity (Fig. 1 C-D, Fig. S3 H).

A key technical challenge in multi-omic assays is cross-library contamination — particularly ATAC fragments appearing in RNA libraries. We found that replacing the Nextera Read 2 adapter with a TruSeq Read 2 adapter in the RNA library preparation effectively resolved this issue for samples with low RNA content. Long-read sequencing confirmed the RNA library structure, with fewer than 0.54% ATAC-derived fragments detected in the RNA library (Table S1).

To benchmark our method, we compared it against 10x Genomics, the current gold standard in single-cell assays. To ensure a fair comparison, we obtained 10x data from ENCODE and published studies using the same K562 cell line, then downsampled all datasets to a uniform read depth per cell (38,375 for ATAC; 69,009 for RNA) based on the sample with the lowest read depth, and called peaks using bulk ATAC-seq from ENCODE. To assess the amount of useful information captured by each assay, we first quantified the number of unique nuclear fragments in peaks per cell. PIP-ATAC-seq performed comparably to 10x ATAC-seq, while PIP-multiome-seq showed lower numbers relative to 10x Multiome-seq. However, the 10x Multiome-seq sample exhibited noticeably lower TSS enrichment, suggesting it may have been over-transposed (Fig. 1F–J). Importantly, both PIP-ATAC-seq and PIP-Multiome-seq displayed higher TSS enrichment than their 10x counterparts, indicating greater specificity and reduced background signal.

To further assess library complexity, we simulated the total number of unique fragments at infinite sequencing depth; both PIP-ATAC-seq and PIP-Multiome-seq were comparable to 10x (Fig. 1E). On the RNA-seq side, PIP-Multiome-seq yielded similar median gene counts per cell compared to 10x Multiome-seq (Fig. 1H–I). Furthermore, the number of unique fragments and detected genes for PIP-ATAC-seq and PIP-Multiome-seq at their original sequencing depths was substantially higher than values reported for other open-source methods based on benchmarking in literature ^7,8^.

We then visually examined bulk ATAC-seq signal tracks for our methods. The PIP-ATAC-seq and PIP-Multiome-seq derived tracks were highly concordant with bulk ATAC-seq from ENCODE and pseudo-bulk tracks from 10x scATAC-seq (Fig. 1J). The tracks also demonstrated high sensitivity, with minor peaks clearly distinguishable against minimal background noise. For the bulk RNA-seq track comparison, we generated a standard PIP-seq RNA library from frozen K562 nuclei; its track was nearly identical to that of our Multiome-seq, suggesting that the additional steps do not significantly degrade RNA-seq quality. Taken together, the distinct signal tracks for each modality indicate no detectable cross-contamination between them. Regarding throughput, 10x can process up to 10,000 cells per lane, whereas PIP-seq can handle between 2,000 and 100,000 cells per sample depending on the kit size. Despite comparable performance, PIP-seq can be 10 times less expensive than 10x, particularly when using this large-scale kit (Fig. 1E).

Finally, we applied PIP-Multiome-seq to frozen mouse brain tissue to assess whether distinct cell types could be identified from a complex tissue sample. We generated two biological replicates and obtained 11,829 nuclei passing QC. The ATAC-seq library yielded a median of 23,055 unique fragments per cell with a TSS enrichment score of 13.1, while the RNA-seq library detected an average of 15,177 transcripts and 4,573 genes per cell (Table S1).To identify cell types, we separately visualized the RNA and ATAC data using uniform manifold approximation and projection (UMAP) in two-dimensional space. Unsupervised clustering revealed 16 distinct cell types, encompassing most major cell types found in the mouse frontal cortex. Notably, the ATAC-seq clustering results were highly concordant with those from RNA-seq, consistent with the established role of chromatin accessibility in regulating gene expression (Fig. 2A–B). Clustering results were also reproducible across both replicates for RNA and ATAC-seq, indicating minimal technical variation (Fig. S4A–B).

**Fig. 2.**
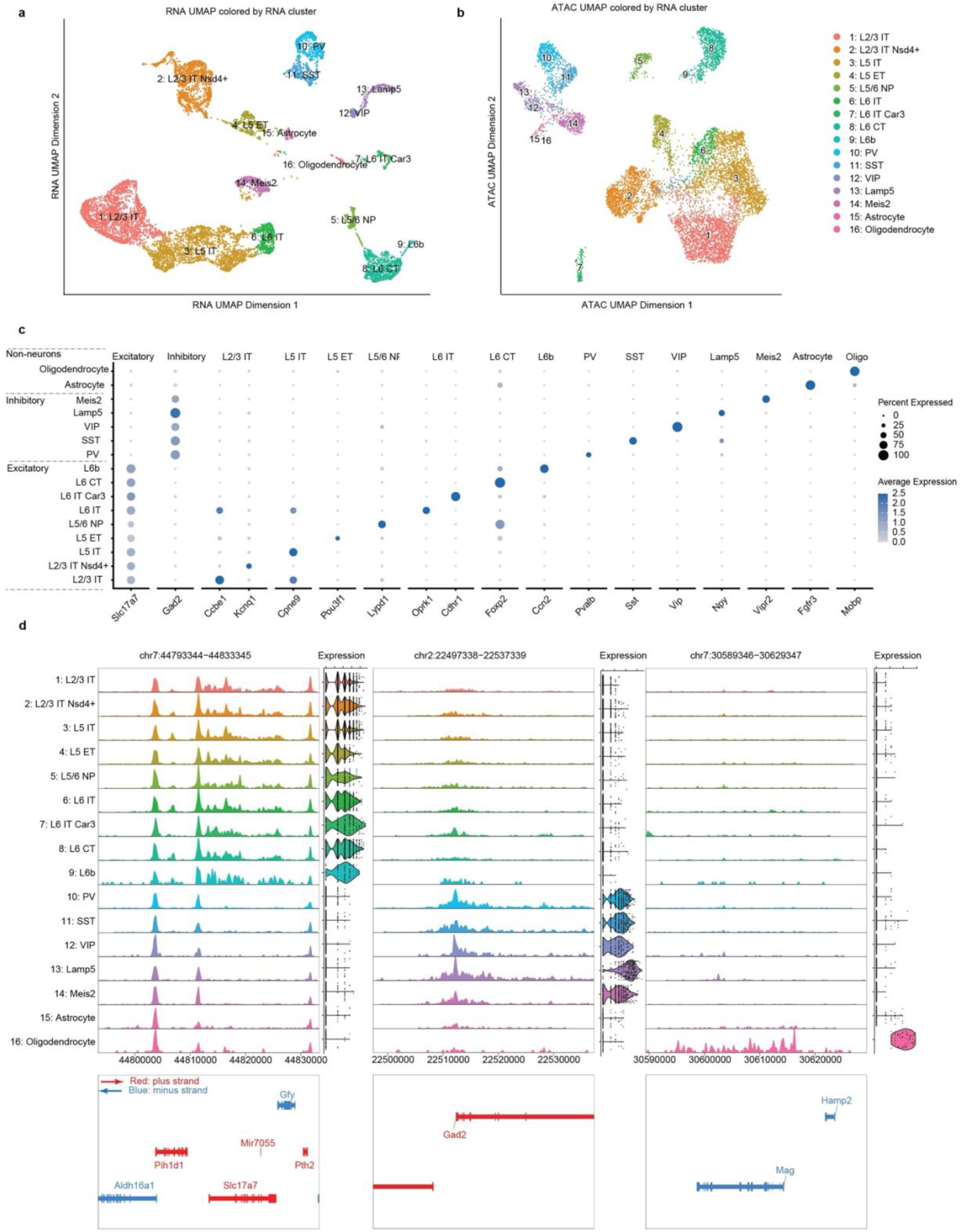
Simultaneous measurement of chromatin accessibility and gene expression in mouse frontal cortex using PIP-Multiome-seq. **a**, UMAP visualization of RNA-based clusters. **b**, UMAP visualization of ATAC-based clusters, colored by RNA cluster identity. **c**, Dot plot showing marker gene expression levels and the percentage of cells expressing each marker gene across clusters annotated by RNA identity. **d**, Pseudo-bulk ATAC-seq tracks at marker gene loci for each cell cluster annotated by RNA identity, with corresponding gene expression shown as violin plots on the side.

To annotate the clusters, we combined automated annotation against a reference dataset, differential gene expression analysis to identify top marker genes, and manual inspection of classical marker expression. This approach identified nine subtypes of excitatory neurons (Slc17a7+) spanning different cortical layers, five subtypes of inhibitory neurons (Gad2+), and two non-neuronal cell types: astrocytes marked by Fgfr3 and oligodendrocytes marked by Mobp (Fig. 2C). Pseudo-bulk ATAC-seq tracks from each cluster exhibited distinct chromatin accessibility signatures around marker gene loci, consistent with their corresponding gene expression patterns (Fig. 2D).

In sum, particle-templated Instant partitioning is a promising strategy for high quality, large scale, low-cost single cell sequencing. Here, we present its first application to chromatin accessibility and multiomics. These applications highlight its potential use as a platform for further method development. For instance, single-cell CUT&TAG, single-cell DNA methylation sequencing, and single-cell footprinting may also be portable to this platform, further increasing the toolkit of single cell molecular characterization methods.

**Fig. S1.**
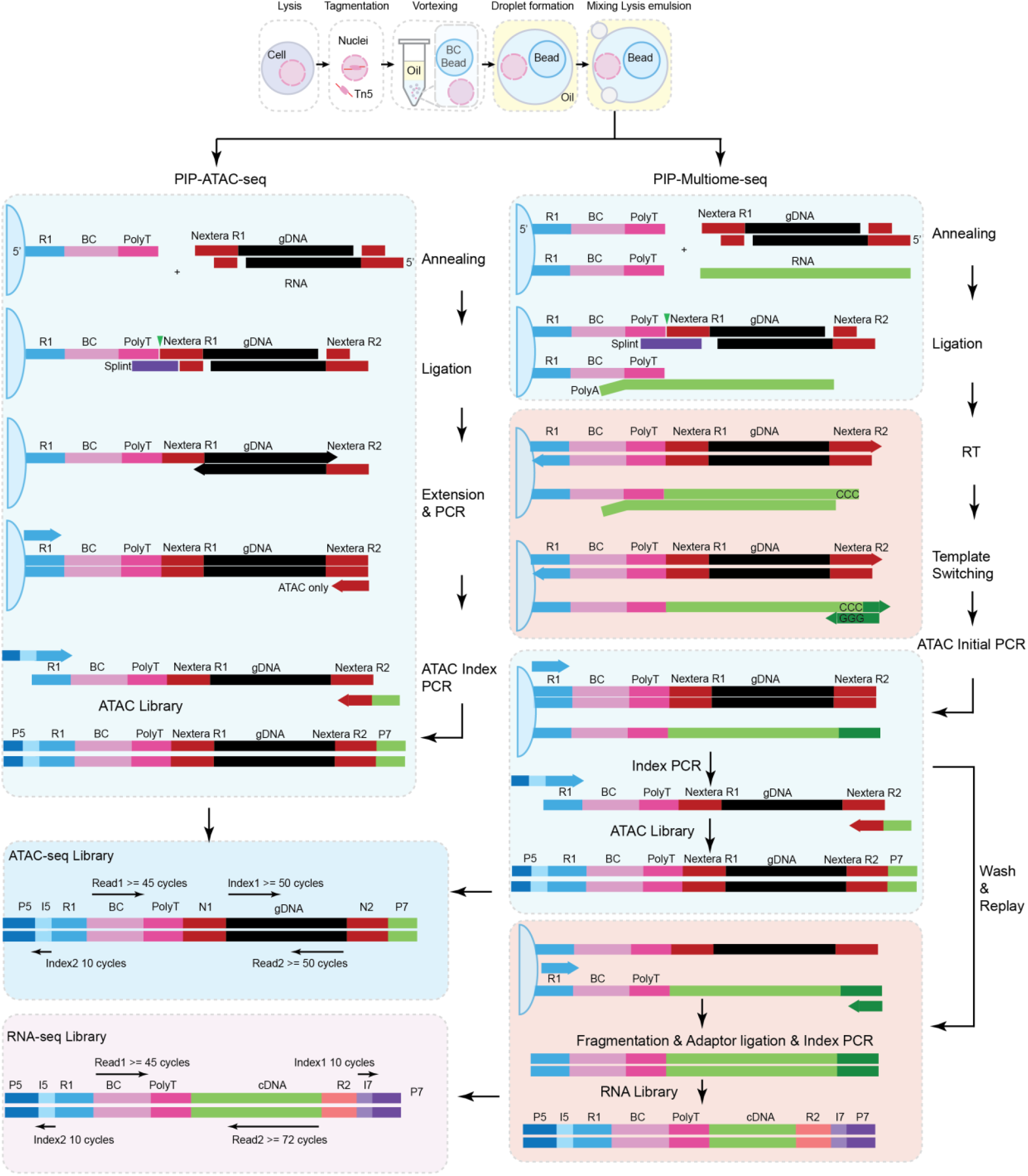
Detailed workflows of PIP-ATAC-seq and PIP-Multiome-seq. The workflows for each method are shown separately.

**Fig. S2.**
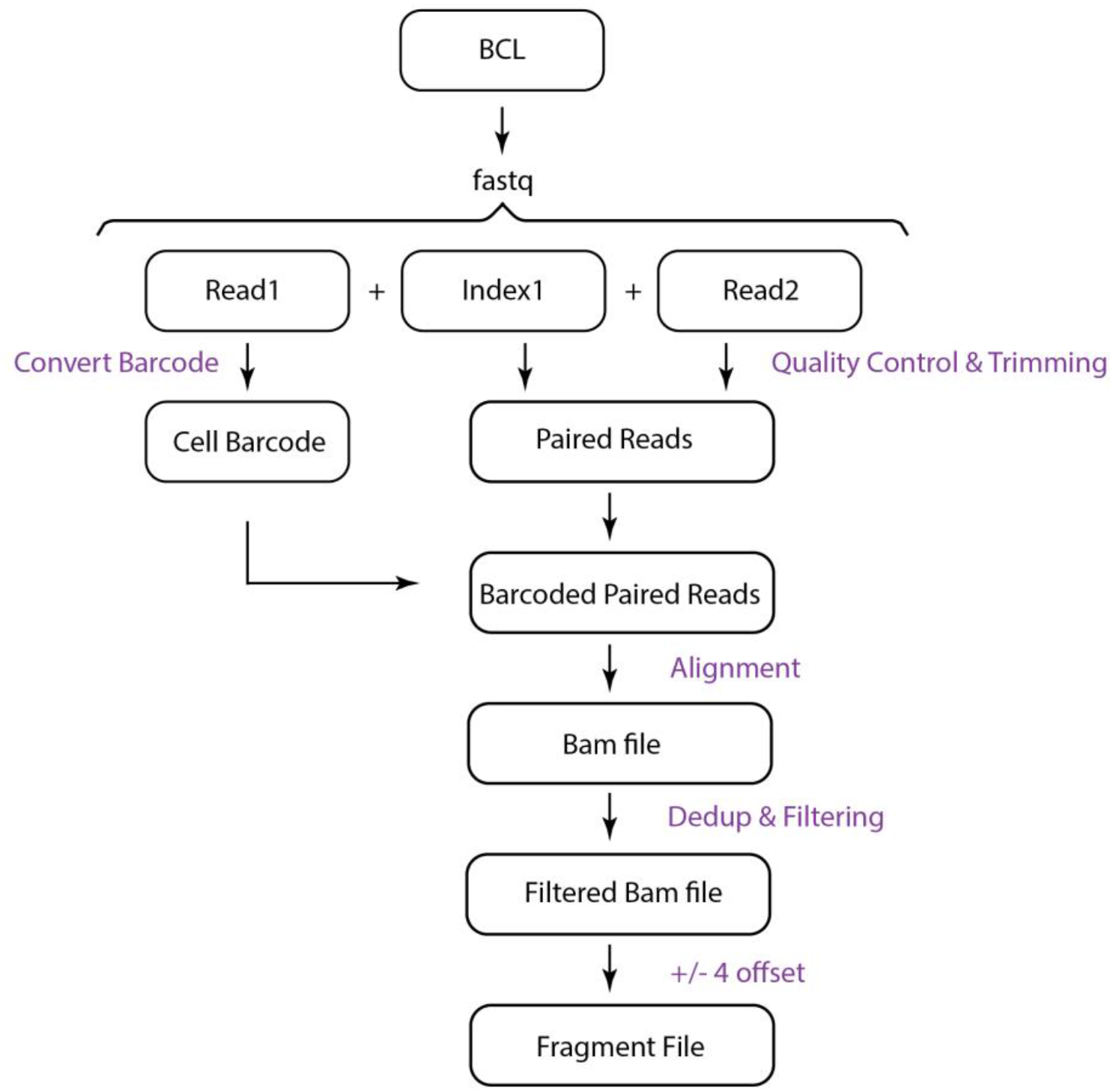
A pre-processing snakemake pipeline to generate ATAC-seq fragment files.

**Fig. S3.**
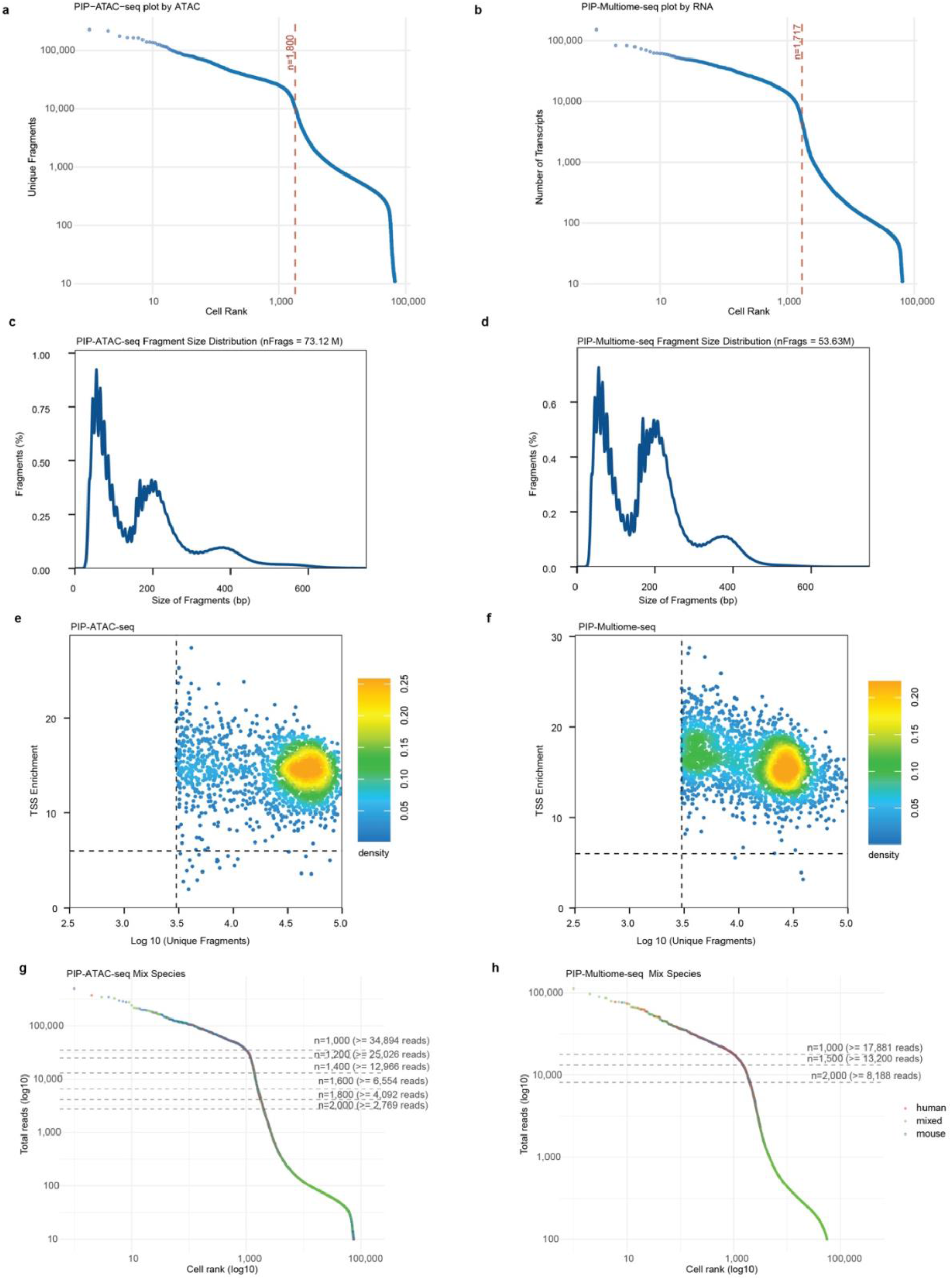
PIP-ATAC-seq and PIP-Multiome-seq data quality for cell lines and species mixing experiments. **a–b**, Elbow plots for selecting the number of cells for downstream analysis: PIP-ATAC-seq (**a**) and PIP-Multiome-seq (**b**). **c–d**, ATAC-seq fragment size distribution for PIP-ATAC-seq (**c**) and PIP-Multiome-seq (**d**). **e–f**, TSS enrichment versus unique fragments plots for visualizing data quality: PIP-ATAC-seq (**e**) and PIP-Multiome-seq (**f**). **g–h**, Elbow plots for species mixing experiments: PIP-ATAC-seq (**g**) and PIP-Multiome-seq (**h**). Cells with more than 90% human reads are shown in red; cells with more than 90% mouse reads are shown in blue; remaining cells are shown in green and classified as mixed.

**Fig. S4.**
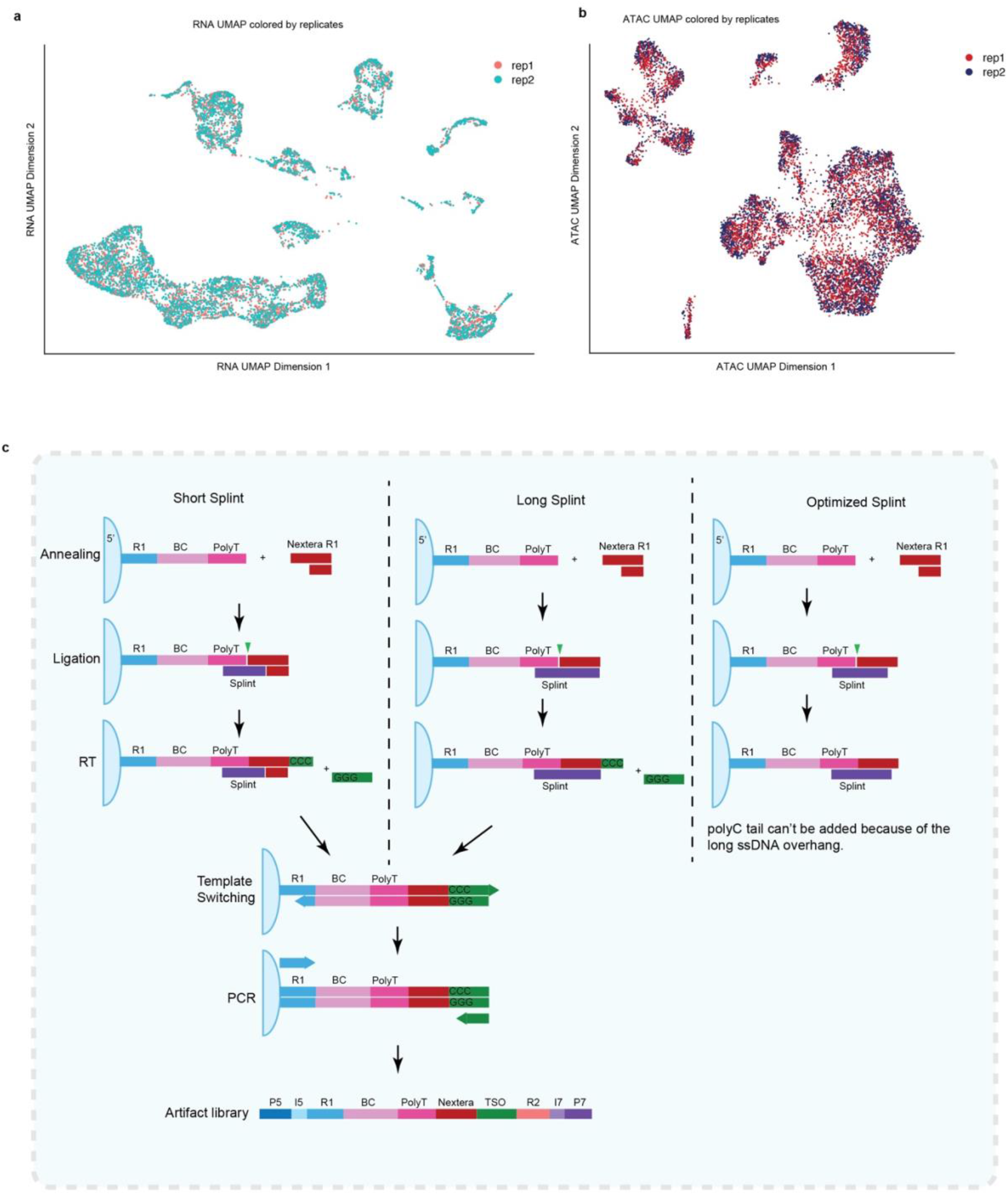
PIP-Multiome-seq mouse brain data quality and splint length optimization. **a–b**, UMAP visualizations colored by replicate identity: RNA-based clusters (**a**) and ATAC-based clusters (**b**). **c**, Splint artifacts can be eliminated by optimizing splint length.

## Methods

### Cell Culture

Human K562 cells were obtained from ATCC and cultured in IMDM (ATCC no. 30-2005) supplemented with 10% FBS and penicillin/streptomycin. Mouse 8024T cells were cultured in DMEM supplemented with 10% FBS and penicillin/streptomycin. All cells were maintained at 37°C with 5% CO_2_. Cells were dissociated using TrypLE and cryopreserved in Bambanker freezing medium (Bulldog Bio, cat. no. BB05) for use in single-cell assays.

### Brain Nuclei Isolation

Fresh-frozen mouse brain tissue from C57BL/6J mice was purchased from Jackson Laboratory. The frontal cortex was dissected and processed for nuclei isolation following a previously described protocol (https://dx.doi.org/10.17504/protocols.io.kxygxmr34l8j/v2).

### PIP-ATAC-seq

See Supplementary Note S1 for a detailed step-by-step protocol.

### PIP-Multiome-seq

See Supplementary Note S2 for a detailed step-by-step protocol. Briefly, the multiome version is a further refined protocol built upon the ATAC-seq version with additional optimizations. Specifically, the length of the splint oligo was optimized to reduce artifact formation introduced during the reverse transcription step. (Fig. S4 B).

### Mixed-Species Experiment

Frozen K562 and 8024T cells were thawed at 37°C. Adherent 8024T cells were passed through a 10 µm cell strainer to reduce clumping. Cells were counted using an automated cell counter in freezing medium. Equal numbers of cells from each species were combined and processed together for nuclei isolation, followed by library preparation for either PIP-ATAC-seq or PIP-Multiome-seq.

### PIP-seq

Frozen K562 cells were thawed and processed for nuclei isolation using the same protocol as PIP-Multiome-seq. A total of 5,000 nuclei were used to prepare standard PIP-seq libraries using the PIP-seq T2 kit following the manufacturer’s nuclei protocol (Fluent Biosciences).

### Sequencing

RNA-seq libraries from PIP-Multiome-seq were sequenced on a NovaSeq X Plus using 2 × 151 bp paired-end reads with a 10 bp index, performed by Novogene. ATAC-seq libraries were sequenced either on a NextSeq 550 using a 150-cycle kit (R1: 45 bp, R2: 50 bp, Index1: 50 bp, Index2: 10 bp) or on a NovaSeq X Plus using a 200-cycle kit (R1: 45 bp, R2: 75 bp, Index1: 75 bp, Index2: 10 bp). Custom sequencing primers were used for ATAC-seq library sequencing as described in Supplementary Notes S1 and S2. Standard PIP-seq libraries were sequenced on a MiSeq using a 150-cycle kit.

### Verification of RNA-seq Library Purity

RNA-seq libraries from PIP-Multiome-seq were submitted to Plasmidsaurus for long-read sequencing. Partial Nextera Read 1 and its reverse complement sequences (TCGTCGGCAGCGTCAGATGTGTATA and TATACACATCTGACGCTGCCGACGA) were used to identify ATAC-seq-derived fragments contaminating the RNA-seq library.

### PIP-ATAC-seq and PIP-Multiome-seq ATAC Data Pre-processing

We developed a Snakemake pipeline enabling parallelized data processing to convert raw FASTQ files to fragment files for downstream analysis (https://github.com/GreenleafLab/PIP-ATAC-Pipeline). The pipeline consists of the following steps (Fig S2):

Quality control. Raw reads were quality-trimmed and adapter sequences were removed using fastp (v1.0.1)^9^. R1 reads containing cell barcodes were processed independently, while I1 and R2 reads containing genomic sequences were processed as paired files. Quality reports were generated for each sample.

Cell barcode extraction. Cell barcodes were extracted from R1 reads using PIPseeker (v3.3.0) with the --chemistry V setting. Barcodes were incorporated into the read headers of the corresponding I1 and R2 reads using a custom Python script that built an in-memory lookup table mapping each R1 read to its 16 bp cell barcode. The barcode was embedded as a CB: tag in the read header to enable barcode-aware downstream processing. Read headers were also cleaned of extraneous nucleotide sequences appended during sequencing.

Alignment and filtering. Quality-controlled I1 and R2 reads (with embedded cell barcodes) were aligned to the reference genome using Bowtie2 (v2.5.4)^10^ in --very-sensitive mode with a maximum insert size of 2,000 bp. Aligned reads were converted to BAM format and processed with SAMtools (v1.20)^11^, including read name sorting, mate-pair fixing, coordinate sorting, and barcode-aware PCR duplicate marking using samtools markdup. Duplicate-marked reads were filtered to retain only properly paired alignments with a minimum mapping quality of MAPQ ≥ 30.

Fragment file generation. Filtered BAM files were converted to fragment files using a custom Python script implemented with pysam (v0.23.3). For each properly paired read, Tn5 transposase insertion sites were identified by applying a +4/−4 bp offset correction to read start positions, which resolves the 1 bp mismatch introduced by the conventional +4/−5 bp offset approach and improves base pair-resolution analysis^3^. Fragment coordinates (chromosome, start, end, and cell barcode) were output in BED format, coordinate-sorted with GNU sort, compressed with bgzip, and indexed with tabix for efficient random-access queries.

Computational infrastructure. All pipeline steps were executed on an HPC cluster via SLURM job scheduling, managed through Snakemake’s SLURM executor plugin. Conda environments were used to ensure software reproducibility.

### PIP-seq and PIP-Multiome-seq RNA Data Pre-processing

Raw FASTQ files (R1 and R2) were processed using PIPseeker (v3.3.0) with the --chemistry V setting. The same reference files used for ATAC-seq alignment were used for RNA-seq. Human data were aligned to the hg38 analysis set without alternative contigs (GCA_000001405.15_GRCh38_no_alt_plus_hs38d1_analysis_set.fna). The GENCODE v49 annotation file was used for human samples, with chromosome names edited to match the reference and filtered to retain only protein-coding, lncRNA, and immunoglobulin genes as recommended by 10x Genomics. Mouse data were aligned to mm39, with the GENCODE vM38 annotation file filtered using the same criteria. Both filtered and unfiltered count matrices were retained for downstream analysis.

### PIP-ATAC-seq and PIP-Multiome-seq QC and Filtering

For PIP-ATAC-seq, fragment files were used to generate read count versus cell rank plots, and real cells were distinguished from background using the elbow point. Fragment files were further analyzed using ArchR (v1.0.3)^12^ with parameters minTSS = 6 and minFrags = 3,000 to remove low-quality cells and quantify unique nuclear fragments and TSS enrichment scores. Only cells passing both filters were retained for downstream analysis. For PIP-Multiome-seq, cell numbers were first estimated from an elbow plot generated by PIPseeker based on RNA-seq data. Cells were further filtered based on the number of unique transcripts, number of detected genes, percentage of mitochondrial reads, TSS enrichment score, and number of unique ATAC fragments. All per-sample filtering thresholds are summarized in Table S1.

### Mixed-Species Analysis

For ATAC-seq, sequenced reads were aligned to a barnyard genome combining hg38 and mm39 using the PIP-ATAC-Pipeline. Real cells were first distinguished from background using an elbow plot, and cells with more than 90% of reads mapping to a single species were assigned to that species; otherwise, they were classified as multiplets. For RNA-seq, FASTQ files were aligned to a barnyard reference (hg38 + mm39 genome; GENCODE v49 + vM38 annotation) using PIPseeker with the --run-barnyard setting. We observed that PIPseeker tended to over-assign reads to three hg38 genes (ENSG00000280441, ENSG00000293668, ENSG00000285708) located within ENCODE blacklist regions; reads mapping to these genes were therefore excluded. Species assignment was based on a threshold of >85% of reads mapping to a single species as reported by PIPseeker.

### Brain Data Analysis

RNA-seq data from QC-filtered cells were used for doublet detection with DoubletFinder (v2.0.6), applied independently to each replicate. For each sample, raw RNA counts were normalized using SCTransform with mitochondrial read percentage regressed out, implemented in Seurat (v5.4.0)^13^. Principal component analysis (PCA) was performed using the top 30 components, followed by UMAP embedding, neighborhood graph construction, and Louvain clustering (resolution = 0.5). The optimal neighborhood size parameter (pK) was determined empirically via a parameter sweep (paramSweep, summarizeSweep). Doublets were identified using DoubletFinder with parameters pN = 0.25, pK = 0.005, and an expected doublet rate of 8%, and predicted doublets were removed prior to downstream analysis.

Singlet-filtered replicates were merged into a single Seurat object and re-normalized using SCTransform (regressing out percent mitochondrial reads), followed by PCA and UMAP embedding using the top 40 principal components. Shared nearest-neighbor graphs were constructed using 40 PCs, and cells were clustered using the Louvain algorithm at resolution 0.8. Clusters were examined following automated cell type annotation by Azimuth (v0.5.0) using the Mouse Motor Cortex reference (mousecortexref). Clusters exhibiting biologically implausible co-expression of markers from distinct cell types — suggestive of residual doublets — or lacking clear cell type-defining marker genes were removed if their cell count was very low or were subclustered and only biologically interpretable subclusters were retained. This served as an additional QC step. After removing low-quality clusters, the process was repeated from unnormalized RNA counts until no further low-quality clusters were identified. Final cluster annotations were based on both automated annotation and canonical marker gene expression.

ATAC-seq data from retained cells were processed using ArchR. Dimensionality reduction was performed using iterative Latent Semantic Indexing (LSI) on the 500 bp tile matrix, retaining the top 25,000 variable features across 2 iterations with 30 LSI dimensions. Cells were clustered using the Seurat-based Louvain algorithm (resolution = 0.8) applied to the LSI embedding. UMAP visualization was generated using 40 neighbors, a minimum distance of 0.4, and cosine distance metric.

### Benchmarking Data Analysis

Raw FASTQ files from 10x Genomics ATAC-seq datasets were downloaded from ENCODE (ENCSR308ZGJ, ENCSR332SXG, ENCSR217VXJ), and 10x Multiome-seq data were downloaded from ArrayExpress (ERR9847049, ERR9847050). The same genome and annotation files used for PIP-ATAC-seq were used to build reference files for 10x data analysis. 10x ATAC-seq and Multiome-seq data were processed using Cell Ranger ATAC (v2.2.0)^14^ and Cell Ranger ARC (v2.1.0), respectively. For 10x ATAC-seq, real cells were identified using an elbow plot and filtered based on TSS enrichment and unique fragment counts. For 10x Multiome-seq, real cells were identified using an elbow plot based on RNA-seq and filtered based on number of genes, number of UMIs, percentage of mitochondrial reads, TSS enrichment, and unique fragment counts. For cross-platform ATAC-seq comparisons, raw FASTQ files were downsampled to match the sequencing depth per cell of the lowest-depth sample prior to reprocessing. TSS enrichment scores and unique fragment counts were assessed in ArchR using cells identified at the original sequencing depth, as fewer cells are typically recovered after downsampling. Peaks were called using MACS3 (v3.0.4)^15^ on a bulk ATAC-seq sample from ENCODE (ENCLB918NXF) and used for all samples. RNA-seq data were similarly compared at 69,009 reads per cell. Detailed statistics are provided in Table S2.

### Estimation of Library Complexity

Aligned BAM files retaining PCR duplicates were filtered by cell barcode to retain only reads from real cells identified at the original sequencing depth using a python script. Library complexity at saturation depth was estimated using the lc_extrap function in Preseq (v3.2.0)^16^ with -B -P for paired end bam file.

## Author Contributions

Z.L. and W.J.G. conceived of the study and Z.L. developed the methods. Z.L. performed the experiments and analyzed the data. Z.L. wrote the analysis pipeline. Z.L. and W.J.G. wrote the manuscript. W.J.G. supervised the work.

## Acknowledgements

We thank Betty B. Liu and Samuel H. Kim for providing the unloaded Tn5. We thank all members of the Greenleaf Lab for helpful discussions. This work is supported by US National Institutes of Health (U19AI057266, R01HG013317, 5R01HL171611, DP1HG013599, R01 NS128028-01) and Arc Institute

