## Supplementary material for "A Simple, Cost-Effective, High-Throughput Method for Measuring Chromatin Accessibility and Gene Expression in Single Nuclei": Note S1

#### Greenleaf Lab PIP-ATAC-Seq Protocol V1

##### Materials

| Item | Vendor | Catalog |
| --- | --- | --- |
| T7 DNA ligase | NEB | M0318L |
| NEBNext Ultra II Q5 Master Mix | NEB | M0544L |
| Digitonin | Promega | G9441 |
| NP-40 | Thermo Fisher Scientific | 85124 |
| Tween 20 | Sigma-Aldrich | P1379-100ML |
| BSA | Sigma | 126609-10gm |
| Protease Inhibitor Cocktail | Sigma-Aldrich | 11873580001 |
| 1M Tris-HCl pH 7.5 | Thermo Fisher Scientific | 15567027 |
| 1M Magnesium acetate | Thermo Fisher Scientific | J60041.AE |
| 1M MgCl2 | Thermo Fisher Scientific | AM9530G |
| 5M Potassium acetate | Sigma | 95843-100ML-F |
| 0.2M Tris Acetate pH 7.8 | Bioworld | 40120265-1 |
| Dimethylformamide (DMF) | Thermo Fisher Scientific | 20673 |
| SPRIselect Reagent Kit | Beckman Coulter | B23318 |
| DNA LoBind Tubes | Eppendorf | 22431048 |
| Disposable Hemocytometer | Fisher Scientific | NC1211379 |
| Ethidium Homodimer-1 (EthD-1) | Thermo Fisher Scientific | E1169 |

### Digitonin is supplied at 2% in DMSO. Dilute 1:1 with water to make a 1% solution to prevent freezing. Can be kept at -20°C for up to 6 months.
### Tween-20 is 10%. Store at 4°C.
### NP40 is 10%. Store at 4°C.

##### Kits

| Item | Vendor | Catalog |
| --- | --- | --- |
| Illumina Single Cell 3' RNA Capture, T2 | Illumina | 20132790 |

### This protocol has been tested for T2 kit.

**Buffers**

#### RSB

| Reagent | Final Concentration | Stock Concentration | Amount to Add (µl) |
|---|---|---|---|
| Tris-HCl pH 7.4 | 10.3 mM | 1 M | 515 |
| NaCl | 10.3 mM | 4 M | 103 |
| MgCl2 | 3.09 mM | 1 M | 155 |
| $H_2O$ | - | - | 49227 |

### Stable at RT.

#### PBS Wash Buffer

| Reagent | Final Conc. | Stock Conc. | 1X (µl) | 2.2X | 3.3X | 4.4X | 8.8X |
|---|---|---|---|---|---|---|---|
| BSA | 0.50% | 25.00% | 20 | 44 | 66 | 88 | 176 |
| PBS | - | - | 980 | 2156 | 3234 | 4312 | 8624 |
| Total | | | 1000 | 2200 | 3300 | 4400 | 8800 |

### Make fresh but can be store at 4C

#### Lysis Buffer

| Reagent | Final Conc. | Stock Conc. | 1X (µl) | 2.2X | 3.3X | 4.4X | 8.8X |
|---|---|---|---|---|---|---|---|
| Tween-20 | 0.10% | 10% | 1 | 2.2 | 3.3 | 4.4 | 8.8 |
| NP-40 | 0.10% | 10% | 1 | 2.2 | 3.3 | 4.4 | 8.8 |
| Digitonin | 0.01% | 1% | 1 | 2.2 | 3.3 | 4.4 | 8.8 |
| RSB | - | - | 97 | 213.4 | 320.1 | 426.8 | 853.6 |
| Total | | | 100 | 220 | 330 | 440 | 880 |

### Make it fresh.

#### Wash Buffer

| Reagent | Final Conc. | Stock Conc. | 1X (µl) | 2.2X | 3.3X | 4.4X | 8.8X |
|---|---|---|---|---|---|---|---|
| Tween-20 | 0.10% | 10% | 10 | 22 | 33 | 44 | 176 |
| BSA | 0.50% | 25.00% | 20 | 44 | 66 | 88 | 352 |
| RSB | - | - | 970 | 2134 | 3201 | 4268 | 17072 |
| Total | | | 1000 | 2200 | 3300 | 4400 | 17600 |

### Make it fresh.

##### 2X Tagmentation Buffer

| Reagent | Final Conc. | Stock Conc. | Amount to Add (µl) |
|---|---|---|---|
| Tris-acetate | 66mM | 0.2 M | 330.0 |
| K-acetate | 132mM | 5 M | 26.4 |
| Mg-acetate | 20mM | 1 M | 20.0 |
| DMF | 32% | 100% | 320.0 |
| BSA | 2.00% | 25.00% | 80.0 |
| $H_2O$ | - | - | 223.6 |

### Can be store at -20°C for a few months.

##### 1X Tagmentation Buffer

| Reagent | Final Conc. | Stock Conc. | 1X(µl) | 2X | 3.3X | 4.4X | 8.8X |
|---|---|---|---|---|---|---|---|
| 2X TB | 1X | 2X | 50 | 55.0 | 82.5 | 110.0 | 220.0 |
| Tween-20 | 0.10% | 10% | 1 | 1.1 | 1.7 | 2.2 | 4.4 |
| Digitonin | 0.01% | 1% | 1 | 1.1 | 1.7 | 2.2 | 4.4 |
| PIC | 0.4X | 50X | 0.8 | 0.9 | 1.3 | 1.8 | 3.5 |
| H2O | - | - | 29.2 | 32.1 | 48.2 | 64.2 | 128.5 |

### Make it fresh.
### Prepare more for 1X to ensure accurate pipetting.
### The amount of Tn5 used per reaction should be adjusted based on its activity and concentration. We use 4µl based on the quality of our in-house purified tn5.

##### TD Buffer

| Reagent | Final Conc. | Stock Conc. | Amount to Add (µl) |
|---|---|---|---|
| Tris-HCl pH 7.6 | 20 mM | 1 M | 40 |
| MgCl2 | 20 mM | 1 M | 40 |
| DMF | 50% | 100% | 500 |
| H2O | - | - | 460 |

##### Oligos

| Name | Concentration |
|---|---|
| Annealed Tn5 adapter | 50µM |
| WTA-F+WTA-R | 5µM |
| Splint oligo | 10µM |
| Truseq R1 + Nextera R2 adapter mix | 10µM |

### Table S2 for details.

**Preparation for loaded Tn5**

**Oligo Annealing**

1. Mix the following reagent in a PCR tube. The volume can be scaled up.

| Reagent | Conc. | Volume(µl) |
|---|---|---|
| ME_TS_A_modified | 100µM | 4.5 |
| ME_TS_B | 100µM | 4.5 |
| ME_NTS_modified | 100µM | 9 |
| NaCl | 500mM | 2 |

2. Preheat a PCR machine. Incubate samples with the below program.

| **Temperature ºC** | **Time** |
|---|---|
| 85 | 2 min |
| Slowly cool to 15°C over 40 min | |
| 4 | Hold |

3. It can be stored at -20°C.

**Tagmentation Test for on bench loading**

Ideally, Tn5 should be mixed with annealed oligos at a 1:1 molar ratio. However, protein concentration estimates may not always be accurate, so it is advisable to test several loading ratios spanning the estimated concentration range. Including a commercially pre-loaded Tn5 as a positive control (e.g., TDE1 from Illumina) is also strongly recommended.

1. Mix Tn5 with annealed adaptors at several different ratios. Incubate at room temperature for 1 hour
2. Assemble the following reaction mixture for each condition. Be sure to include a no-Tn5 negative control.

| Reagent | Volume(µl) |
|---|---|
| Loaded Tn5 | 1.5 |
| gDNA | 75ng |
| 2X TD Buffer | 10 |
| $H_2O$ | Up to 20 |

3. Incubate at 55°C for 7min.
4. Purify using a Zymo DNA purification kit.
5. Assess fragment size distribution by TapeStation, Bioanalyzer, or gel electrophoresis.

Select the ratio that yields the most fragmented genomic DNA. It is recommended to test different amounts of loaded Tn5 using bulk ATAC-seq. The optimal amount should generate libraries with clear nucleosome periodicity and without large peaks several thousand base pairs in size. Use this same amount of Tn5 per sample when proceeding to the single-cell protocol below.

#### Tn5 on bench loading

Mix Tn5 with annealed adaptors at the optimal ratio determined above. Incubate at room temperature for 1 hour. Loaded Tn5 can be stored at −20°C for up to several months.

### This protocol has been validated for use with live and frozen cells. Bambanker freezing medium is recommended for cell preservation.

### Nuclei isolation can be done according to a demonstrated protocol from 10X. “Nuclei Isolation for Single Cell Multiome ATAC + Gene Expression Sequencing (CG000365)”.

### Nuclei quality is the key. It’s recommended to first test the nuclei quality after isolation using fluorescence dye (EthD-1) for new sample types. Extend time in the lysis buffer if not all the nuclei can be stained positive. Reduce it if nuclei are damaged etc.

Nuclei isolation for cell lines (>100,000 cells)

1. Thaw cells rapidly in a 37 °C water bath if frozen. Remove immediately once thawed.
2. Count cells and transfer 100,000–1,000,000 cells into a 1.5 mL DNA LoBind tube.
3. For frozen samples, slowly add 1 mL culture medium dropwise to the tube.
4. If significant clumping is present, optionally filter the suspension through an appropriate cell strainer.
5. Centrifuge at 300 × g for 5 min and carefully remove the supernatant.
   Tip: Use a P1000 followed by a P200 pipette to minimize sample loss throughout the procedure.
6. Resuspend the pellet in 1 mL PBS wash buffer. Centrifuge at 300 × g for 5 min and remove the supernatant.
7. Resuspend the pellet in 100 µL ATAC lysis buffer and incubate on ice for 3 min.
8. Add 1 mL ATAC wash buffer. Centrifuge at 500 × g for 4 min and remove the supernatant.
9. Add 1 mL ATAC wash buffer. Determine nuclei concentration using an automated cell counter. Centrifuge at 500 × g for 4 min and remove the supernatant.
10. Based on the concentration from Step 9, resuspend nuclei in ATAC wash buffer to a final concentration of 10,000 nuclei/µL.

Nuclei isolation for cell lines when input is limited (<100,000 cells)

1. Thaw cells rapidly in a 37 °C water bath if frozen. Remove immediately once thawed.
2. Count cells and transfer 60,000 cells into a 1.5 mL DNA LoBind tube.
3. For frozen samples, slowly add 1 mL culture medium dropwise to the tube.
4. Centrifuge at 300 × g for 5 min and carefully remove the supernatant.
   Tip: Use a P1000 followed by a P200 pipette to minimize sample loss throughout the procedure.
5. Resuspend the pellet in 1 mL PBS wash buffer. Centrifuge at 300 × g for 5 min and remove the supernatant.
6. Resuspend the pellet in 50 µL ATAC lysis buffer and incubate on ice for 3 min.
7. Add 1 mL ATAC wash buffer, then centrifuge at 500 × g for 5 min. Remove the supernatant.
8. Resuspend in 5 µL ATAC wash buffer. Skip this step if more than 5 µL wash buffer remains after Step 7.

Nuclei isolation for tissue.

Follow the protocol "Isolation of nuclei from frozen tissue for snMultiome, snATAC-seq, snRNA-seq, and other epigenomic/transcriptomic assays V.2", stopping at step 39. Skip the transposition of nuclei section. (dx.doi.org/10.17504/protocols.io.kxygxmr34l8j/v2).

**Transposition**

1. Transfer 4~5 µL of nuclei suspension (50,000 nuclei) into a 1.5 mL DNA LoBind tube. Add 41 µL tagmentation buffer and mix thoroughly.
2. Add 4 µL loaded Tn5 and mix gently but thoroughly.
3. Incubate at 37 °C for 30 min with shaking at 800 rpm.
4. Prepare 3 mL NSB working solution per sample according to the "Prepare Nuclei Suspension" section of the PIP-seq protocol without RNase Inhibitor.
5. Add ~950 µL NSB working solution (from the PIP-seq kit) and mix well by pipetting. Centrifuge at 500 × g for 4 min and carefully remove the supernatant.
6. Resuspend in 1 mL NSB working solution and transfer to a new 1.5 mL DNA LoBind tube. Centrifuge at 500 × g for 4 min and carefully remove the supernatant.
7. Resuspend in 1 mL NSB working solution, centrifuge at 500 × g for 4 min, and carefully remove the supernatant. Perform three washes total.

**T2 Kit**
**Droplet formation**

1. Resuspend nuclei in 40 µL NSB working solution (for the T2 kit).
2. Add 1 µL splint oligo to the sample and mix thoroughly.
3. Optional: Take 3 µl of the nuclei suspension and mix with either 3 µl of trypan blue or 3 µl of diluted EthD-1 (prepared by mixing 1µl 2 mM EthD-1 with 50 µl PBS). Load the mixture onto a Bulldog hemocytometer to count nuclei and assess nuclei quality. Nuclei should appear single, intact, and 100% stained.
4. Generate droplets according to the "Capture and Lysis" section of the PIP-seq protocol. Add 4–5 µL nuclei suspension (depending on concentration) to one tube of PIP-seq beads.
5. Incubate under nuclei lysis conditions. Use a thermomixer with a ThermoTop, without shaking, and set the cooling rate to the minimum. Alternatively, a PIP-seq dry bath may be used, but the yield may be slightly reduced. Overnight incubation is recommended as a stopping point.

**Ligation**

1. Follow the "Isolate mRNA" section from the PIP-seq protocol. The break emulsion and wash the beads before DNA synthesis.
2. Add following ligation reagents into the beads. Mix well by pipetting.

| Reagent | 1X (µl) | 2.2X | 3.3X | 4.4X | 8.8X |
|---|---|---|---|---|---|
| T7 Ligase | 4 | 8.8 | 13.2 | 17.6 | 35.2 |
| 2X quick ligation buffer | 44 | 96.8 | 145.2 | 193.6 | 387.2 |

3. Incubate at 25ºC for 30min.
4. Put the tube in the red rack. Take out the supernatant above the wire line and keep ~39µl.
5. Wash the beads twice with 150µl 0.5X cold wash buffer according to the PIP-seq protocol in PCR strips.

**ATAC-seq library generation**

**Initial Amplification**

1. Perform initial PCR amplification by adding below reagents into beads and mix well.

| Reagent | 1X (μl) | 2.2X | 3.3X | 4.4X | 8.8X |
|---|---|---|---|---|---|
| 5μM of WTA-F +WTA-R | 10 | 22 | 33 | 44 | 88 |
| Ultra II Q5 | 50 | 110 | 165 | 220 | 440 |

2. Amplify for 5 cycles using the below setting.

| Cycle | Temperature ºC | Time (s) |
|---|---|---|
| **Extension** | 72 | 300 |
| **Denature** | 98 | 30 |
| **10X** | 98 | 10 |
| | 68 | 30 |
| | 72 | 30 |
| **Final Extension** | 72 | 30 |
| **Hold** | 4 | - |

3. Take 60μl of supernatant. Save in a new tube.
4. Add 40μl of buffer CE, mix, spin down for 10 seconds, then take out 40μl.
5. Combine the total 100μl of supernatant. Add 100μl of SPRI beads for 1X purification.
6. Incubate for 5 min at room temperature.
7. Put it on a magnetic wait till the solution is clear.
8. Add 200μl of 85% EtOH, wait 30 seconds, remove the EtOH.
9. Repeat step 8.
10. Add 20μl of IDTE buffer incubate for 5 min.
11. Put it on a magnetic wait till the solution is clear. Save the supernatant in a new tube.

**Final Amplification**

1. Add below reagents into 20μl of the library for a total 50μl.

| Reagent | Amount (μl) |
|---|---|
| 2X NEBNext Ultra II Q5 | 25 |
| 10μM Truseq R1 + Nextera R2 adapter mix | 5 |

2. Adjust the cycle number based on sample type (see Example Library section).

| Cycle | Temperature ºC | Time (s) |
|---|---|---|
| **Denature** | 98 | 30 |
| **~10X** | 98 | 10 |
| | 68 | 30 |
| | 72 | 30 |
| **Final Extension** | 72 | 30 |
| **Hold** | 4 | - |

3. 40μl of SPRI beads for a 0.8X purification.
4. Incubate for 5 min at room temperature.
5. Put it on a magnetic wait till the solution is clear.
6. Add 200μl of 85% EtOH, wait 30 seconds, remove the EtOH.
7. Repeat step 8.
8. Add 20μl of IDTE buffer incubate for 5 min.
9. Put it on a magnetic wait till the solution is clear.

10. Quantify supernatant with Qubit and check with Tapestation D5000 / D5000 high sensitive tape. Save it in a 1.5ml DNA lobind tube.

**Example Library**
The following is a representative ATAC-seq library prepared from K562 cells using the T2 kit. Final PCR: 10 cycles. Concentration: 30.7 ng/µl in 20 µl.

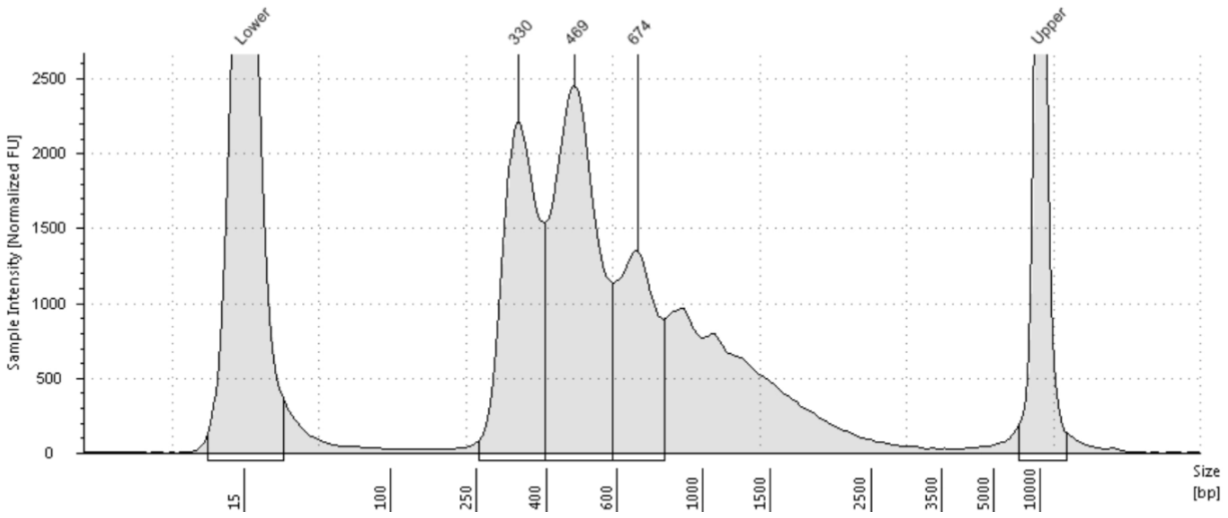


**Sequencing**
The ATAC-seq library will be sequenced using custom sequencing primers.
Read1: >= 45 cycles (Use Truseq Read1 as custom sequencing primer)
Read2: >= 50 cycles (Default Illumina primer)
Index1: >= 50 cycles (Use Nextera Read1 as custom sequencing primer)
Index2: 10 cycles (Use Truseq Read1 RC as custom sequencing primer)
#Skip index2 primers on machines using forward strand workflow.

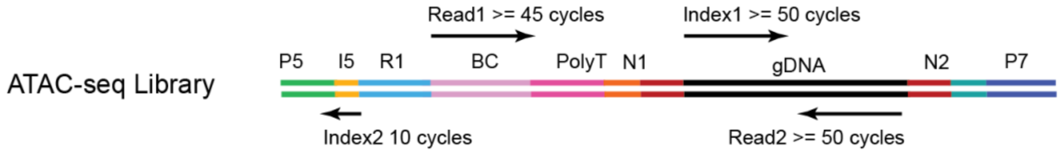
