## Supplementary material for "A Simple, Cost-Effective, High-Throughput Method for Measuring Chromatin Accessibility and Gene Expression in Single Nuclei": Note S2

### Greenleaf lab PIP-Multiome-seq Protocol V1.4

#### Materials

| Item | Vendor | Catalog |
|---|---|---|
| T7 DNA ligase | NEB | M0318L |
| Thermolabile Exonuclease I | NEB | M0568L |
| Deoxynucleotide (dNTP) Solution Mix | NEB | N0447S |
| Q5U Hot Start High-Fidelity DNA Polymerase | NEB | M0515L |
| NEBNext Ultra II Q5 Master Mix | NEB | M0544L |
| Digitonin | Promega | G9441 |
| NP-40 | Thermo Fisher Scientific | 85124 |
| Tween 20 | Sigma-Aldrich | P1379-100ML |
| BSA | Sigma | 126609-10gm |
| Protease Inhibitor Cocktail | Sigma-Aldrich | 11873580001 |
| RNase Inhibitor | Qiagen | Y9240L |
| Protector RNase Inhibitor | Sigma | 3335402001 |
| 1M Tris-HCl pH 7.5 | Thermo Fisher Scientific | 15567027 |
| 1M Magnesium acetate | Thermo Fisher Scientific | J60041.AE |
| 1M MgCl2 | Thermo Fisher Scientific | AM9530G |
| 5M Potassium acetate | Sigma | 95843-100ML-F |
| 0.2M Tris Acetate pH 7.8 | Bioworld | 40120265-1 |
| Dimethylformamide (DMF) | Thermo Fisher Scientific | 20673 |
| SPRIselect Reagent Kit | Beckman Coulter | B23318 |
| DNA LoBind Tubes | Eppendorf | 22431048 |
| Disposable Hemocytometer | Fisher Scientific | NC1211379 |
| Ethidium Homodimer-1 (EthD-1) | Thermo Fisher Scientific | E1169 |

### Digitonin is supplied at 2% in DMSO. Dilute 1:1 with water to make a 1% solution to prevent freezing. Can be kept at -20°C for up to 6 months.
### Tween-20 is 10%. Store at 4°C.
### NP40 is 10%. Store at 4°C.
### Protector RNase Inhibitor is used in all buffers except the tagmentation buffer, which uses Qiagen RNase Inhibitor.

#### Kits

| Item | Vendor | Catalog |
|---|---|---|
| Illumina Single Cell 3' RNA Prep, T2 | Illumina | 20135689 |
| Illumina Single Cell 3' RNA Prep, T10 | Illumina | 20135691 |

### This protocol has been tested for T2 and T10 kits.

#### Buffers

##### RSB

| Reagent | Final Concentration | Stock Concentration | Amount to Add (µl) |
|---|---|---|---|
| Tris-HCl pH 7.4 | 10.3 mM | 1 M | 515 |
| NaCl | 10.3 mM | 5 M | 103 |
| MgCl2 | 3.09 mM | 1 M | 155 |
| $H_2O$ | - | - | 49227 |

### Stable at RT.

##### PBS Wash Buffer

| Reagent | Final Conc. | Stock Conc. | 1X (µl) | 2.2X | 3.3X | 4.4X | 8.8X |
|---|---|---|---|---|---|---|---|
| BSA | 0.50% | 25.00% | 20 | 44 | 66 | 88 | 176 |
| PBS | - | - | 980 | 2156 | 3234 | 4312 | 8624 |
| Total | | | 1000 | 2200 | 3300 | 4400 | 8800 |

### Make fresh but can be store at 4°C for weeks.

##### Lysis Buffer

| Reagent | Final Conc. | Stock Conc. | 1X (µl) | 2.2X | 3.3X | 4.4X | 8.8X |
|---|---|---|---|---|---|---|---|
| Tween-20 | 0.10% | 10% | 1 | 2.2 | 3.3 | 4.4 | 8.8 |
| NP-40 | 0.10% | 10% | 1 | 2.2 | 3.3 | 4.4 | 8.8 |
| Digitonin | 0.01% | 1% | 1 | 2.2 | 3.3 | 4.4 | 8.8 |
| RSB | - | - | 97 | 213.4 | 320.1 | 426.8 | 853.6 |
| Total | | | 100 | 220 | 330 | 440 | 880 |
| RNase Inhibitor | 0.8X | 40X | 2 | 4.4 | 6.6 | 8.8 | 17.6 |

### Make it fresh.

##### Wash Buffer

| Reagent | Final Conc. | Stock Conc. | 1X (µl) | 2.2X | 3.3X | 4.4X | 8.8X |
|---|---|---|---|---|---|---|---|
| Tween-20 | 0.10% | 10% | 22 | 44 | 66 | 88 | 176 |
| BSA | 0.50% | 25.00% | 44 | 88 | 132 | 176 | 352 |
| RSB | - | - | 2134 | 4268 | 6402 | 8536 | 17072 |
| Total | | | 2200 | 4400 | 6600 | 8800 | 17600 |
| RNase Inhibitor | | | 5.5 | 11 | 16.5 | 22 | 44 |

### Make it fresh.

##### 2X Tagmentation Buffer

| Reagent | Final Conc. | Stock Conc. | Amount to Add (µl) |
|---|---|---|---|
| Tris-acetate | 66mM | 0.2 M | 330.0 |
| K-acetate | 132mM | 5 M | 26.4 |
| Mg-acetate | 20mM | 1 M | 20.0 |
| DMF | 32% | 100% | 320.0 |
| BSA | 2.00% | 25.00% | 80.0 |
| $H_2O$ | - | - | 223.6 |

### Can be store at -20°C for a few months.

##### 1X Tagmentation Buffer

| Reagent | Final Conc. | Stock Conc. | 1X(µl) | 2X | 3.3X | 4.4X | 8.8X |
|---|---|---|---|---|---|---|---|
| 2X TB | 1X | 2X | 50 | 55.0 | 82.5 | 110.0 | 220.0 |
| Tween-20 | 0.10% | 10% | 1 | 1.1 | 1.7 | 2.2 | 4.4 |
| Digitonin | 0.01% | 1% | 1 | 1.1 | 1.7 | 2.2 | 4.4 |
| PIC | 0.4X | 50X | 0.8 | 0.9 | 1.3 | 1.8 | 3.5 |
| Rnase In | 0.8X | 40X | 2 | 2.2 | 3.3 | 4.4 | 8.8 |
| H2O | - | - | 27.2 | 29.9 | 44.9 | 59.8 | 119.7 |

### Make it fresh.
### Prepare more for 1X to ensure accurate pipetting.
### Make sure to use Qiagen RNase Inhibitor Y9240L because it doesn't inhibit Tn5.
### The amount of Tn5 used per reaction should be adjusted based on its activity and concentration. We use 4µl based on the quality of our in-house purified tn5.

##### TD Buffer

| Reagent | Final Conc. | Stock Conc. | Amount to Add (µl) |
|---|---|---|---|
| Tris-HCl pH 7.6 | 20 mM | 1 M | 40 |
| MgCl2 | 20 mM | 1 M | 40 |
| DMF | 50% | 100% | 500 |
| H2O | - | - | 460 |

### Can be store at -20°C for a few months.

##### Oligos

| Name | Concentration |
|---|---|
| Annealed Tn5 adapter | 50µM |
| Biotin TSO | 50µM |
| WTA-F | 10µM |
| PCR_blk+ | 100µM |
| Splint oligo | 10µM |
| Annealed TruSeq R2 ligation adapter | 50µM |
| Truseq R1 + Truseq R2 adapter mix | 10µM |
| Truseq R1 + Nextera R2 adapter mix | 10µM |

### Table S2 for details.

**Preparation for loaded Tn5**
**Oligo Annealing**

1. Mix the following reagent in a PCR tube. The volume can be scaled up.

| Reagent | Conc. | Volume(μl) |
|---|---|---|
| ME_TS_A_modified | 100μM | 4.5 |
| ME_TS_B | 100μM | 4.5 |
| ME_NTS_modified | 100μM | 9 |
| NaCl | 500mM | 2 |

2. Preheat a PCR machine. Incubate samples with the below program.

| Temperature ºC | Time |
|---|---|
| 85 | 2 min |
| Slowly cool to 15°C over 40 min | |
| 4 | Hold |

3. It can be stored at -20°C.

It is preferable to load the oligo onto Tn5 on column, which requires starting from the protein purification step. Alternatively, it can be loaded easily on the bench using the protocol listed below.

**Tagmentation Test for on bench loading**

Ideally, Tn5 should be mixed with annealed oligos at a 1:1 molar ratio. However, protein concentration estimates may not always be accurate, so it is advisable to test several loading ratios spanning the estimated concentration range. Including a commercially pre-loaded Tn5 as a positive control (e.g., TDE1 from Illumina) is also strongly recommended.

1. Mix Tn5 with annealed adaptors at several different ratios. Incubate at room temperature for 1 hour
2. Assemble the following reaction mixture for each condition. Be sure to include a no-Tn5 negative control.

| Reagent | Volume(μl) |
|---|---|
| Loaded Tn5 | 1.5 |
| gDNA | 75ng |
| 2X TD Buffer | 10 |
| $H_2O$ | Up to 20 |

3. Incubate at 55°C for 7min.
4. Purify using a Zymo DNA purification kit.
5. Assess fragment size distribution by TapeStation, Bioanalyzer, or gel electrophoresis.

Select the ratio that yields the most fragmented genomic DNA. It is recommended to test different amounts of loaded Tn5 using bulk ATAC-seq. The optimal amount should generate libraries with clear nucleosome periodicity and without large peaks several thousand base pairs in size. Use this same amount of Tn5 per sample when proceeding to the single-cell protocol below.

**Tn5 on bench loading**

Mix Tn5 with annealed adaptors at the optimal ratio determined above. Incubate at room temperature for 1 hour. Loaded Tn5 can be stored at −20°C for up to several months.

### This protocol has been validated for use with live cells, frozen cells, fresh-frozen brain tissue, and cryopreserved PBMCs. Bambanker freezing medium is recommended for cell preservation.

**Ligation**

1. Follow the “Isolate mRNA” section from PIP-seq protocol. The break emulsion and wash the beads before DNA synthesis.
2. Add following ligation reagents into the beads. Mix well by pipetting.

| Reagent | 1X (µl) | 2.2X | 3.3X | 4.4X | 8.8X |
|---|---|---|---|---|---|
| T7 Ligase | 4 | 8.8 | 13.2 | 17.6 | 35.2 |
| RI (protector 40U/µl) | 1 | 2.2 | 3.3 | 4.4 | 8.8 |
| 2X quick ligation buffer | 44 | 96.8 | 145.2 | 193.6 | 387.2 |

3. Incubate at 25ºC for 30min.
4. Put the tube in the red rack. Take out the supernatant above the wire line and keep ~39µl.
5. Wash the beads twice with 150µl 1X cold wash buffer according to the PIP-seq protocol in PCR strips.

**Synthesize cDNA**

1. Proceed to the “Synthesize cDNA” section in PIP-seq protocol.
2. Replace TSO1 in the RT master mix with an equal amount of 50µM Biotin-TSO.
3. Using below setting on a thermomixer instead of the PIP-seq cDNA synthesis program. It can be hold overnight at 4C.

| Temperature ºC | Time (min) |
|---|---|
| 25 | 30 |
| 42 | 90 |
| 60 | 10 |
| 4 | Hold |

**Exonuclease treatment.**

1. Wash the beads with 150µl 1X washing buffer for a total of three washes.
2. Add below reagents into ~39µl of beads and mix well.

| Reagent | 1X (µl) | 2.2X | 3.3X | 4.4X | 8.8X |
|---|---|---|---|---|---|
| 10X NEB buffer 3 | 5 | 11 | 16.5 | 22 | 44 |
| Thermolabile Exo 1 | 2.5 | 5.5 | 8.25 | 11 | 22 |

3. Incubate at 37ºC, 1000rpm for 10min in a thermomixer.
4. Wash the beads four times with 150µl 0.5X wash buffer for a total of four washes. Proceed to the next step quickly.

**ATAC-seq library generation**

**Initial Amplification**

1. Perform initial PCR amplification by adding below reagents into beads and mix well.

| Reagent | 1X (µl) | 2.2X | 3.3X | 4.4X | 8.8X |
|---|---|---|---|---|---|
| 5µM of WTA F | 6 | 13.2 | 19.8 | 26.4 | 52.8 |
| Ultra II Q5 | 0.6 | 1.32 | 1.98 | 2.64 | 5.28 |
| 5X NEBnext Q5 Buffer | 12 | 26.4 | 39.6 | 52.8 | 105.6 |
| 10mM dNTPs | 1.2 | 2.64 | 3.96 | 5.28 | 10.56 |
| H2O | 1.2 | 2.64 | 3.96 | 5.28 | 10.56 |

2. Amplify for 10 cycles using below setting.

| Cycle | Temperature ºC | Time (s) |
|---|---|---|
| Denature | 98 | 30 |
| 10X | 98 | 10 |
| | 67 | 30 |
| | 72 | 30 |
| Final Extension | 72 | 30 |
| Hold | 4 | - |

3. Add 40µl of buffer CE, mix, spin down for 10 seconds.
4. Take 60µl of supernatant. Save in a new tube.
5. Add 60µl of buffer CE, mix, spin down for 10 seconds, then take out 60µl.
6. Combine the total 120µl of supernatant. Add 144µl of SPRI beads for 1.2X purification.
7. Incubate for 5 min at room temperature.
8. Put it on a magnetic wait till the solution is clear.
9. Add 200µl of 85% EtOH, wait 30 seconds, remove the EtOH.
10. Repeat step 8.
11. Add 20µl of IDTE buffer incubate for 5 min.
12. Put it on a magnetic wait till the solution is clear. Save the supernatant in a new tube.

**Final Amplification**

1. Add below reagents into 20µl of the library for a total 50µl.

| Reagent | Amount (µl) |
|---|---|
| 2X NEBNext Ultra II Q5 | 25 |
| 10µM Truseq R1 + Nextera R2 adapter mix | 5 |

2. Adjust the cycle number based on sample type (see Example Library section).

| Cycle | Temperature ºC | Time (s) |
|---|---|---|
| Denature | 98 | 30 |
| ~10X | 98 | 10 |
| | 68 | 30 |
| | 72 | 30 |
| Final Extension | 72 | 30 |
| Hold | 4 | - |

3. 40µl of SPRI beads for a 0.8X purification.
4. Incubate for 5 min at room temperature.
5. Put it on a magnetic wait till the solution is clear.
6. Add 200µl of 85% EtOH, wait 30 seconds, remove the EtOH.
7. Repeat step 8.
8. Add 20µl of IDTE buffer incubate for 5 min.
9. Put it on a magnetic wait till the solution is clear.
10. Quantify supernatant with Qubit and check with Tapestation D5000 / D5000 high sensitive tape. Save it in a 1.5ml DNA lobind tube.

**RNA-seq library generation**

1. Wash beads four times with 150µl 0.5X wash buffer.
2. Follow "Amplify cDNA" to "Clean Up Library" section in the PIP-seq protocol for RNA-seq part with three changes listed below
3. In the "Amplify cDNA" section, add 0.6µl of 100µM PCR blk+ into each sample for cDNA amplification.
4. In the "Ligate Adapters" section, replace the PIP-seq library adapter mix (LAM) with an equal volume of 50µM Annealed TruSeq R2 ligation adapter.
5. In the "Amplify Library" section, add 0.5µl of 100µM PCR blk+ (1µl for low RNA samples such as PBMC) into each sample for index PCR. Replace the Index adapters with 10µM Truseq R1 + Truseq R2 adapter mix.

**T10 Kit**
**Droplet formation**

1. Resuspend nuclei in 8μL NSB working solution (for the T10 kit).
2. Optional: Measure the volume of the nuclei suspension using a P20 pipette. If excess liquid is present, centrifuge at 300 × g for 3 min, then carefully remove the excess supernatant.
3. Add 1 μL splint oligo to the sample and mix thoroughly.
4. Optional: Take 2 μl of the nuclei suspension and mix with either 4 μl of trypan blue or 4 μl of diluted EthD-1 (prepared by mixing 1 μl 2mM EthD-1 with 50 μl PBS). Load the mixture onto a Bulldog hemocytometer to count nuclei and assess nuclei quality. Adjust nuclei concentration as needed by adding more NSB working solution or by centrifuging at 300 × g for 3min and carefully remove excessive supernatant. Nuclei should appear mostly single, intact, and 100% stained. Don't proceed if nuclei are heavily clumped
5. Generate droplets according to the "Capture and Lysis" section of the PIP-seq protocol. Add 5μL nuclei suspension and 1 μL RNase inhibitor (Protector, 40 U/μL) to one tube of PIP-seq beads.
6. Incubate under nuclei lysis conditions. Use a thermomixer with a ThermoTop, without shaking, and set the cooling rate to the minimum. Alternatively, a PIP-seq dry bath may be used, but the yield may be slightly reduced. Overnight incubation is recommended as a stopping point.

**Ligation**

1. Follow the "Isolate mRNA" section from PIP-seq protocol. The break emulsion and wash the beads before DNA synthesis.
2. Add following ligation reagents to the beads in a 0.5ml tube. Mix well by pipetting.

| Reagent | 1X (μl) | 2.2X | 3.3X | 4.4X | 8.8X |
|---|---|---|---|---|---|
| T7 Ligase | 10 | 22 | 33 | 44 | 88 |
| RI (protector 40U/μl) | 2.5 | 5.5 | 8.25 | 11 | 22 |
| 2X quick ligation buffer | 112.5 | 247.5 | 371.25 | 495 | 990 |

3. Incubate at 25ºC for 30min.
4. Take out the supernatant above the 100μl mark.
5. Wash the beads twice with 450μl 1X cold wash buffer according to the PIP-seq protocol in 0.5ml tube.

**Synthesize cDNA**

1. Proceed to the "Synthesize cDNA" section in PIP-seq protocol.
2. Replace TSO1 in the RT master mix with an equal amount of 50μM Biotin-TSO.
3. Using below setting on a thermomixer instead of the PIP-seq cDNA synthesis program. It can be held overnight at 4C.

| Temperature ºC | Time (min) |
|---|---|
| 25 | 30 |
| 42 | 90 |
| 60 | 10 |
| 4 | Hold |

**Exonuclease treatment.**

1. Wash the beads with 450μl 1X washing buffer for a total of three washes.
2. Add below reagents into ~39μl of beads and mix well.

| Reagent | 1X (μl) | 2.2X | 3.3X | 4.4X | 8.8X |
|---|---|---|---|---|---|
| 10X NEB buffer 3 | 12 | 26.4 | 39.6 | 52.8 | 105.6 |
| Thermolabile Exo 1 | 6 | 13.2 | 19.8 | 26.4 | 52.8 |

3. Incubate at 37ºC, 1000rpm for 10min in a thermomixer.
4. Wash the beads four times with 450μl 0.5X wash buffer for a total of four washes. Proceed to the next step quickly.

**ATAC-seq library generation**

**Initial Amplification**

1. Perform initial PCR amplification by adding below reagents to beads and mix well. Divide ~75μl into PCR strips.

| Reagent | 1X (μl) | 2.2X | 3.3X | 4.4X | 8.8X |
|---|---|---|---|---|---|
| 5X NEBnext Q5 Buffer | 30 | 66 | 99 | 132 | 264 |
| 5μM of WTA F | 15 | 33 | 49.5 | 66 | 132 |
| Ultra II Q5 | 1.5 | 3.3 | 4.95 | 6.6 | 13.2 |
| 10 mM dNTPs | 3 | 6.6 | 9.9 | 13.2 | 26.4 |

2. Amplify for 10 cycles using below setting.

| Cycle | Temperature ºC | Time (s) |
|---|---|---|
| Denature | 98 | 30 |
| 10X | 98 | 10 |
| | 67 | 30 |
| | 72 | 30 |
| Final Extension | 72 | 30 |
| Hold | 4 | - |

3. Add 100μl of buffer CE into each tube of PCR strip, mix, combine all the ~350μl solutions into a 1.5ml DNA lobind tube.
4. Spin for 30s, take out 200μl supernatant, save in a new 1.5ml DNA lobind tube.
5. Wash the PCR tube with 125μl of buffer CE to collect leftover beads, then add to the 1.5ml tube containing beads.
6. Spin for 30s, take out 150μl supernatant, combine for a total 350μl of supernatant in the 1.5ml tube.
7. Add 420μl of SPRI beads for 1.2X purification.
8. Incubate for 5 min at room temperature.
9. Put it on a magnetic wait till the solution is clear.
10. Add 1ml of 85% EtOH, wait 30 seconds, remove the EtOH.
11. Repeat step 8.
12. Add 20μl of IDTE buffer incubate for 5 min.
13. Put it on a magnetic wait till the solution is clear. Save the supernatant in a new tube.

**Final Amplification**

1. Add below reagents into 20µl of the library for a total 50µl.

| Reagent | Amount (µl) |
|---|---|
| 2X NEBNext Ultra II Q5 | 25 |
| 10µM Truseq R1 + Nextera R2 adapter mix | 5 |

2. Amplify using the below program. Adjust the cycle number based on sample type (see Example Library section).

| Cycle | Temperature ºC | Time (s) |
|---|---|---|
| Denature | 98 | 30 |
| ~10X | 98 | 10 |
| | 68 | 30 |
| | 72 | 30 |
| Final Extension | 72 | 30 |
| Hold | 4 | - |

3. 40µl of SPRI beads for a 0.8X purification.
4. Incubate for 5 min at room temperature.
5. Put it on a magnetic wait till the solution is clear.
6. Add 200µl of 85% EtOH, wait 30 seconds, remove the EtOH.
7. Repeat step 8.
8. Add 20µl of IDTE buffer incubate for 5 min.
9. Put it on a magnetic wait till the solution is clear.
10. Quantify supernatant with Qubit and check with Tapestation D5000 / D5000 high sensitive tape. Save it in a 1.5ml DNA lobind tube.

**RNA-seq library generation**

1. Wash beads four times with 1ml 0.5X wash buffer.
2. Follow "Amplify cDNA" to "Clean Up Library" section in the PIP-seq protocol for RNA-seq part with three changes listed below
3. In the "Amplify cDNA" section, add 1.4 µl of 100µM PCR blk+ into each sample (1µM final concentration) for cDNA amplification.
4. In the "Ligate Adapters" section, replace the PIP-seq library adapter mix (LAM) with an equal volume of 50µM Annealed TruSeq R2 ligation adapter.
5. In the "Amplify Library" section, add 0.5µl of 100µM PCR blk+ (1µl for low RNA samples such as PBMC) into each sample for index PCR. Replace the Index adapters with 10µM Truseq R1 + Truseq R2 adapter mix.

**Example Library**
The following is a representative ATAC-seq library prepared from K562 cells using the T2 kit. Final PCR: 10 cycles. Concentration: 5.15 ng/µl in 20 µl.

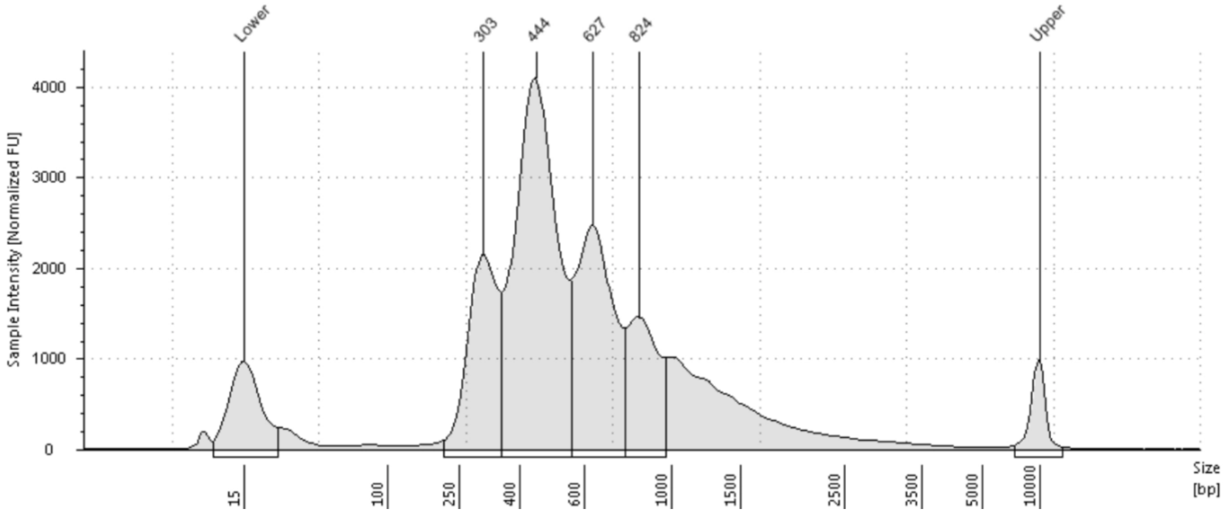


The following is a representative RNA-seq library prepared from K562 cells using the T2 kit. Final PCR: 10 cycles. Concentration: 12.6 ng/µl in 20 µl.

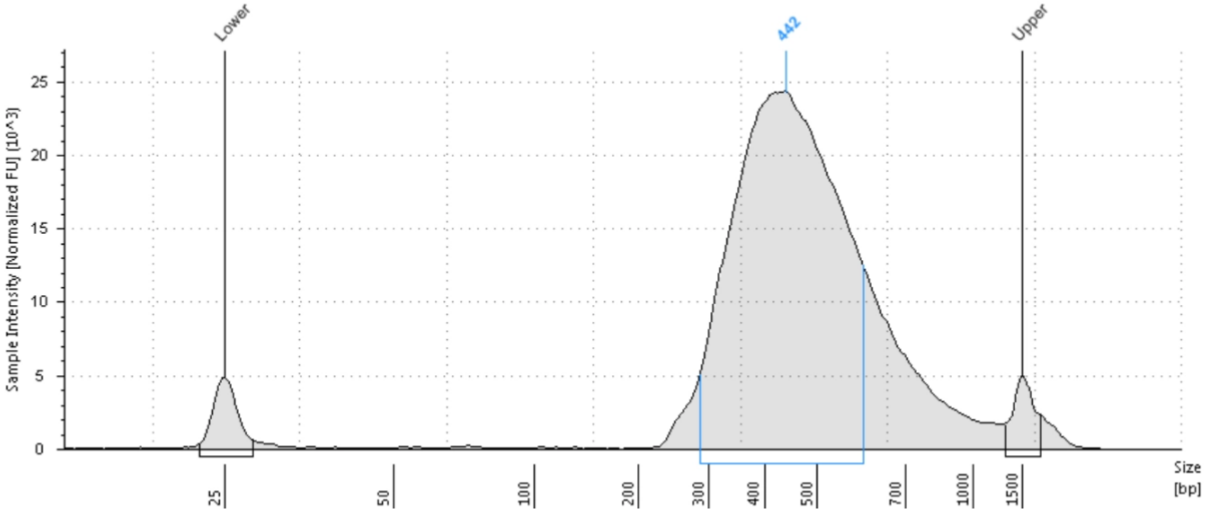


Library prepared from mouse brain using T10 kit.
ATAC-seq: Final PCR: 10 cycles. Concentration: 17.6 ng/µl in 20 µl.
RNA-seq: Final PCR: 10 cycles. Concentration: 8.89 ng/µl in 20 µl.

**Sequencing**
The RNA-seq library can be sequenced according to standard PIP-seq protocol
Read1: >= 45 cycles
Read2: >= 72cycles
Index1: 10 cycles
Index2: 10 cycles

The ATAC-seq library will be sequenced using custom sequencing primers.
Read1: >= 45 cycles (Use Truseq Read1 as custom sequencing primer)
Read2: >= 50 cycles (Default Illumina primer)
Index1: >= 50 cycles (Use Nextera Read1 as custom sequencing primer)
Index2: 10 cycles (Use Truseq Read1 RC as custom sequencing primer)
#Skip index2 primers on machines using forward strand workflow.

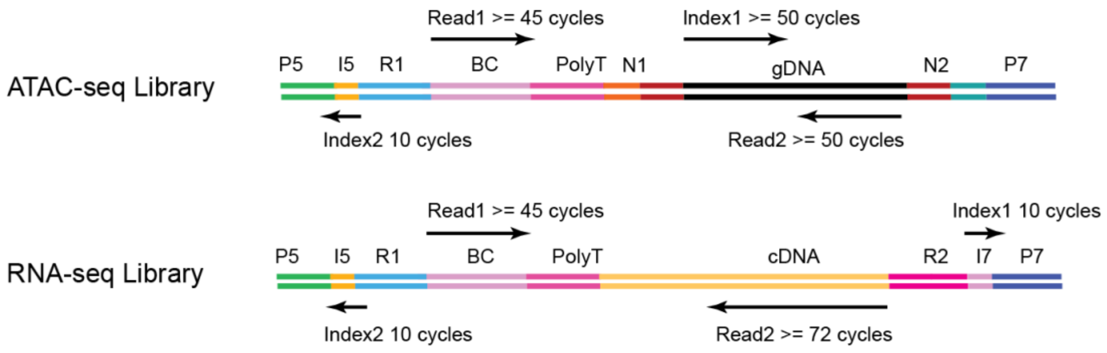
